# Non-Coding and Loss-of-Function Coding Variants in *TET2* are Associated with Multiple Neurodegenerative Diseases

**DOI:** 10.1101/759621

**Authors:** J. Nicholas Cochran, Ethan G. Geier, Luke W. Bonham, J. Scott Newberry, Michelle D. Amaral, Michelle L. Thompson, Brittany N. Lasseigne, Anna M. Karydas, Erik D. Roberson, Gregory M. Cooper, Gil D. Rabinovici, Bruce L. Miller, Richard M. Myers, Jennifer S. Yokoyama

## Abstract

We conducted genome sequencing to search for rare variation contributing to early onset Alzheimer’s disease (EOAD) and frontotemporal dementia (FTD). Discovery analysis was conducted on 493 cases and 671 controls of European ancestry. Burden testing for rare variation associated with disease was conducted using filters based on variant rarity (less than 1 in 10,000 or private), computational prediction of deleteriousness (CADD 10 or 15 thresholds), and molecular function (protein loss-of-function only, coding alteration only, or coding plus non-coding variants in experimentally predicted regulatory regions).

Replication analysis was conducted on 16,871 independent cases and 15,941 independent controls. Rare variants in *TET2* were enriched in the discovery combined EOAD and FTD cohort (*p*=6.5×10^−8^, genome-wide corrected *p*=0.0037). Most of these variants were canonical loss-of-function or non-coding in predicted regulatory regions. This enrichment replicated across several cohorts of AD and FTD (replication only *p*=0.0071). The combined analysis odds ratio was 2.2 (95% CI 1.5–3.2) for AD and FTD. The odds ratio for qualifying non-coding variants considered independently from coding variants was 2.1 (95% CI 1.2–3.9). For loss-of-function variants, the combined odds ratio (for AD, FTD, and amyotrophic lateral sclerosis, which shares clinicopathological overlap with FTD) was 3.2 (95% CI 2.0–5.3). TET2 catalyzes DNA demethylation. Given well-defined changes in DNA methylation that occur during aging, rare variation in *TET2* may confer risk for neurodegeneration by altering the homeostasis of key aging-related processes. Additionally, our study emphasizes the relevance of non-coding variation in genetic studies of complex disease.

## INTRODUCTION

Neurodegeneration with a clinical onset prior to the age of 65 can be devastating for patients, their families, and caregivers, imposing financial burden and hardship during a period of life when individuals are often most productive^1^. Early-onset neurodegenerative diseases such as early-onset Alzheimer’s disease (EOAD) and frontotemporal dementia (FTD) are typically thought of as disease forms with highly penetrant genetic contributions, and indeed both can result from Mendelian pathogenic mutations (with Mendelian causes more common in FTD)^2^. However, these diseases exhibit a high degree of heritability that remains unexplained by currently known genetic contributors^3; 4^. This suggests that additional genetic factors likely contribute to disease but have not yet been identified. Despite attempts at genome-wide association study (GWAS) of relatively sizeable cohorts, only modest association signals have been identified for FTD^5^ and one form of EOAD, posterior cortical atrophy^6^. In contrast, sequencing studies have been successful at identifying more moderately to highly penetrant contributions to disease by examining rare variation. Successes in Alzheimer’s disease (AD) include *ABCA7*, *SORL1*, and *TREM2* (reviewed in ^7^). Similar successes for the amyotrophic lateral sclerosis (ALS)-FTD spectrum include *TBK1*^8^, *MFSD8*^9^, *DPP6*, *UNC13A*, and *HLA-DQA2*^10^. Despite these successes, the rarity of these diseases along with the high cost of sequencing studies has resulted in limited sample sizes of patient cohorts. Furthermore, prior studies have focused on coding regions of the genome, leaving non-coding regions largely unexplored for their contribution to disease risk.

Here we leveraged a large cohort of 683 patients, many of which have an early age of disease onset (<65), and 856 healthy adult controls (with no known neurological abnormalities) that have undergone genome sequencing to probe both coding and non-coding rare and predicted deleterious variants across the genome for association with disease risk. We assessed variant associations between EOAD and FTD vs. controls both separately and together (all cases versus controls), with the hypothesis that genetic pleiotropy—where variation in a single gene associates with multiple, different phenotypes— may play a role, as previously described for neurodegenerative diseases^11–15^.

## METHODS

### Sample selection

The majority of cases were selected from the University of California, San Francisco (UCSF) Memory and Aging Center with an intentional selection of early-onset cases when possible to maximize the likelihood of identifying genetic contributors, along with healthy older adult controls (a total of 664 cases and 102 controls, with 71 of these cases previously described in ^9^). All UCSF cases and controls were clinically assessed using methods described in ^9^. This cohort was intentionally depleted of cases with known Mendelian variants associated with neurodegenerative diseases, and any cases with known Mendelian variants identified after genome sequencing were excluded (see **Results**). A small number of samples (19 cases and 21 controls) were obtained from the University of Alabama at Birmingham (UAB) from an expert clinician who employed the same diagnostic procedures (case studies described in ^16^). The resulting cohort was enriched for early-onset cases with a median age of presentation of 60 (range 45–84) for AD and 66 (range 29–89) for FTD. Additional neurologically healthy controls sequenced at HudsonAlpha were also included from two cohorts: a healthy aging control set from the National Institute of Mental Health (NIMH) (132 controls) and healthy unaffected parents from a childhood disease study where *de novo* mutations are the most common cause of disease^17^, making these parents reasonably representative population controls (601 controls). All participants or their surrogates provided written informed consent to participate in this study and the institutional review boards at each site approved all aspects of the study.

### Genome sequencing

The majority of genome sequencing was performed at the HudsonAlpha Institute for Biotechnology on the Illumina HiSeq X platform (1,468 samples from UCSF, UAB, NIMH, and HudsonAlpha), while a small subset was sequenced at the New York Genome Center, also on the HiSeq X platform (71 samples from UCSF, described previously in ^9^). Mean depth was 34X with an average of 92% of bases covered at 20X. Sequencing libraries at HudsonAlpha were prepared by Covaris shearing, end repair, adapter ligation, and PCR using standard protocols. Library concentrations were normalized using KAPA qPCR prior to sequencing. All variants meeting either Mendelian diagnostic criteria or variants in top hits from the discovery cohort were validated by Sanger sequencing.

### Data quality control

All sequencing reads from both sequencing centers were aligned to the hg19 reference genome with bwa-0.7.12^18^. BAMs were sorted and duplicates were marked with Sambamba 0.5.4^19^. Indels were realigned, bases were recalibrated, and gVCFs were generated with GATK 3.3^20^. Variants were called across all samples in a single batch with GATK 3.8 using the -newQual flag to minimize false negative singleton calls. The VCF was quality filtered with a genotype level requirement for 95% of sites to have a minimum GQ of 20 and DP of 10 (applied using VCFtools 0.1.15^21^), and a variant level filter of VQSLOD > - 3. The small number of missing genotypes remaining after that quality filtering step were assumed to be reference (filled with bcftools 1.6-19^22^) in order to avoid errors in downstream processing using the package GEMINI 0.20.2^23^ which adds missing genotypes to non-reference counts with its burden function. Goleft indexcov 0.1.17^24^ was used for sex checks and samples failing sex checks were excluded. KING 2.1.2^25^ was used to check for familial relationships and related individuals (up to 4^th^ degree relatives using IBD segment analysis) were excluded. Ancestry was elucidated by both principal component analysis using plink 1.9^26^ compared to 1000 genomes data^27^ (using common variation overlapping with 1000 genomes calls) and analysis using ADMIXTURE 1.3.0^28^ (**Supplemental Figure 1**), and only samples from the largest cluster (European ancestry) were retained for discovery analysis to minimize potential confounding population effects.

**Figure 1:**
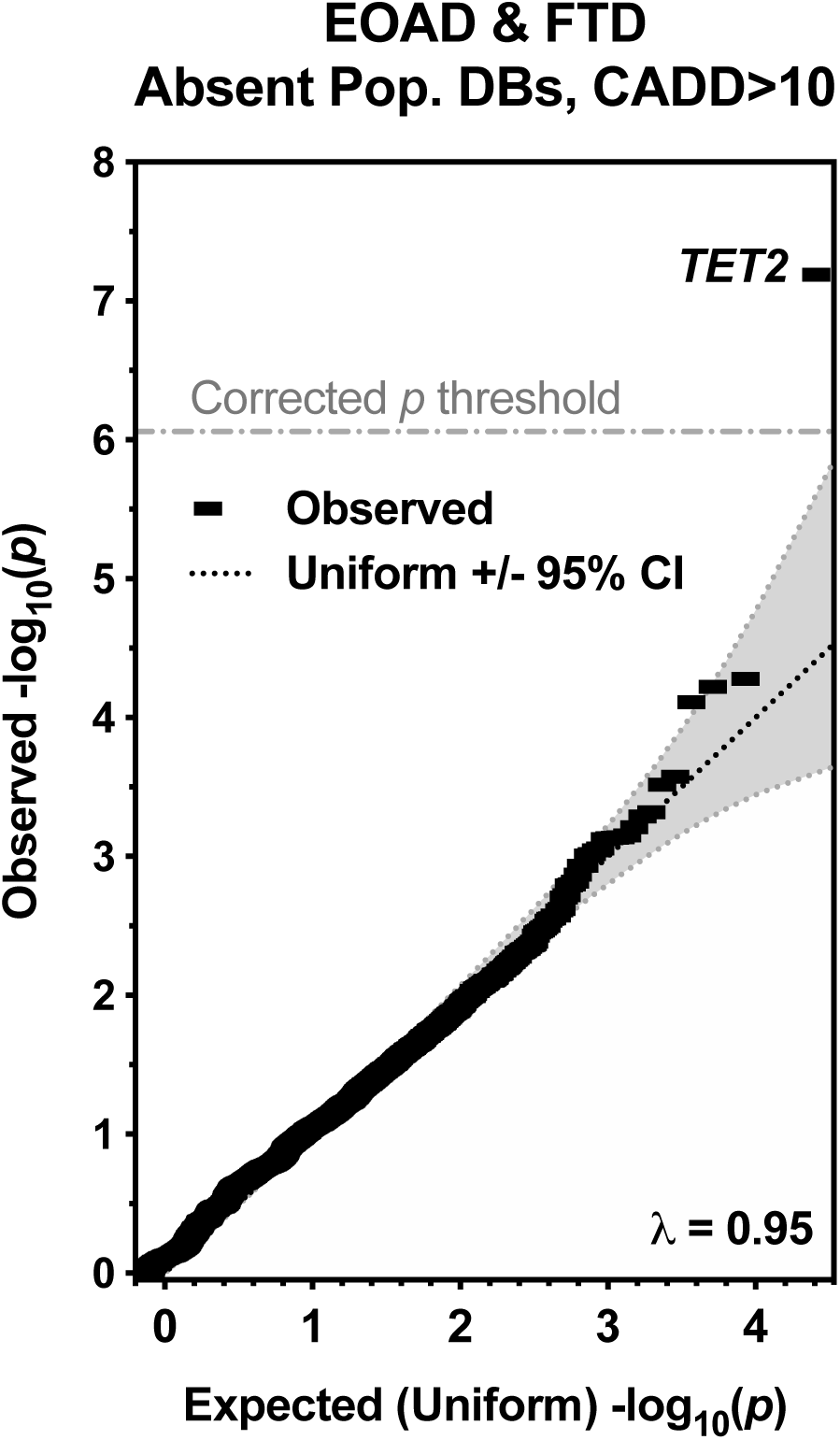
QQ plot of *p*-values from the discovery burden analysis of EOAD and FTD cases versus controls and Private, CADD > 10 Variants. *TET2* is the top and only hit reaching statistical significance (corrected *p*<0.05). No genomic inflation was observed (λ = 0.95). The uniform distribution and theoretical 95% confidence interval based on a beta distribution is shown. Note that, in addition to passing the correction threshold, *TET2* also falls well outside of theoretical random *p*-value distributions.

### Annotation, filtering, and burden analysis

In order to facilitate annotation and burden analysis, multi-allelic sites were split using Vt^29^. All variants were annotated with CADD v1.3^30^, including all indels. SnpEff 4.3s^31^ was used to annotate with the GRCh37.75 gene model. Population database frequency annotations included 1000 genomes phase 3, TOPMed Bravo^32^ (lifted over from hg38 to hg19 using CrossMap 0.2.7^33^), and several population database sets annotated using WGSA 0.7^34^ including ExAC^35^, gnomAD^36^, ESP, and UK10K. Variants were also annotated with dbSNP release 151^37^. A final important annotation set was the union of regions called by GenoSkyline-Plus^38^ as potential regulatory regions. GenoSkyline-Plus incorporates chromatin marks, DNA accessibility, RNA-seq, and DNA methylation to predict function. All tracks derived from direct human tissue sources were included (sources propagated in culture were excluded), with a total of 50 of 66 tissue and cell types described in Table S2 of^38^ used for annotation (see **Supplemental Table 1** for included epigenome tracks in the union).

Variants were filtered using SnpSift 4.3s. In addition to the quality filters described, variants were further filtered by local and population frequency, predicted deleteriousness (CADD v1.3), and segmentation for function. To enrich for rare variation, variants were pre-filtered for a maximum minor allele count of 3 (approximately 0.1% local allele frequency), and a maximum allele frequency of 1 in 10,000 in any population included in the aforementioned population databases. In addition, non-coding variants were more strictly filtered to only variants present in a GenoSkyline-Plus qualifying region as described above and required to be absent from dbSNP 151.

From the initial pre-filtered file, we conducted further filtering to arrive at nine total filter conditions. First, we evaluated variants meeting either: 1 in 10,000 population cutoffs and CADD score greater than 10 or 15, or: private variation and CADD score greater than 10 or 15; for a total of four conditions that include non-coding variants. We also confined to coding variants with the same allele frequency combinations and CADD cutoffs listed for four total coding-only conditions. For canonical loss-of-function, we only considered the base 1 in 10,000 allele frequency requirement and CADD 10 cutoff for a total of one canonical loss-of-function condition (also note that all canonical loss-of-function variants meeting these criteria were included in the other eight filter conditions regardless of allele frequency or CADD cutoff given the known deleteriousness of canonical loss-of-function variants). We note that these are extensively overlapping test sets (See **Supplemental Figure 2** for correlations), and thus often yield similar results. For example, all conditions constrained to private variation will be a subset of matched conditions with 1 in 10,000 population frequency cutoffs, and all coding-only conditions are a subset of the conditions that allow rare non-coding variation.

**Figure 2:**
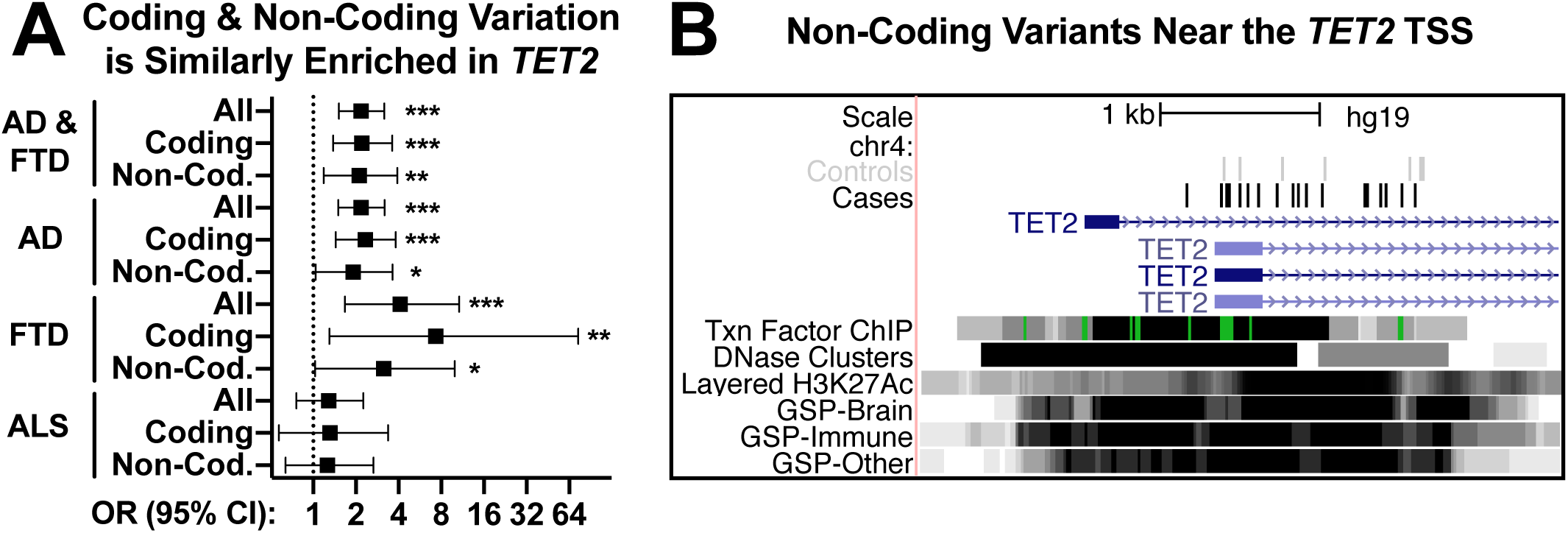
Qualifying non-coding variation in *TET2* is enriched at a similar level as coding variation and occurs in key predicted functional regulatory regions. **A.** Odds ratios are shown for combined analyses (cohorts described in **Table 1**). Breaking out coding and non-coding variation reveals similar effect sizes and *p*-values. * indicates *p*<0.05, ** indicates *p*<0.01, and *** indicates *p*<0.001 by Fisher’s exact test. **B.** Non-coding variants near the *TET2* transcription start site (hg19 chr4:106,066,000-106,070,000) serve as an example of variant enrichment in key regions predicted to have regulatory function. GSP indicates GenoSkyline-Plus regions.

A VCF for each of the nine filtering conditions was loaded into a GEMINI 0.20.2 database^23^, which was used to aggregate counts of variants for each individual by gene. By default, GEMINI is constrained to coding variation, so GEMINI python scripts were edited to allow for counting of variants in non-coding regions as well. Variants upstream or downstream (within 5kb, the SnpEff default) were also assigned to their adjacent genes. The number of qualifying individuals was the final count unit, where one or more qualifying variants in a gene for a given individual resulted in that individual having a qualifying count for that gene (i.e., if an individual had two qualifying variants, they would still only be counted once to account for the possibility of a recessive model of inheritance or negligibility of the 2^nd^ variant if on the same allele). Individuals with more than three qualifying variants in a gene were not counted as qualifying because an excess of rare and predicted damaging variants in a single gene may be indicative of a sequencing or variant-calling error.

### Burden analysis statistics

In order to assess the effect of covariates for the discovery set as well as any replication sets where the necessary covariate data were available, we tested using SKAT 1.3.2.1^39^ using the adaptive efficient re-sampling method corrected for sex, number of *APOE* ε4 alleles, the first four principal components from common variant PCA, and ancestral proportions from ADMIXTURE with k=5. Statistical significance was set at a corrected *p*-value < 0.05. Because of the three main clusters of filter conditions corresponding to case-control test sets (EOAD vs. control, FTD vs. control, or all cases vs. control) (**Supplemental Figure 2**), we applied a correction factor of three to all protein coding genes in hg19 put forth by the HUGO Gene Nomenclature Committee (19,118 genes, **Supplemental Table 2**) for a correction factor for *p* values in the discovery analysis of 57,354. In order to allow for use of replication cohorts where covariate data was not available, we also utilized a two-sided Fisher’s exact test. SKAT and Fisher’s tests were highly correlated (Pearson’s r=0.76 of log transformed *p* values).

### Clinical rate of progression analysis

Our study utilized genetic and longitudinal clinical data from the Alzheimer’s Disease Neuroimaging Initiative (ADNI) to study the clinical profiles and progression of *TET2* rare variant carriers. ADNI is a multi-center prospective longitudinal cohort study created to study the genetic, clinical, and imaging correlates of Alzheimer’s disease^40–42^, and ADNI cases are present in the Alzheimer’s Disease Sequencing Project (ADSP) replication cohort. Every study participant undergoes a thorough assessment that includes clinical characteristics, cognitive testing, and genetic sequencing. Participants were diagnosed as either normal controls (CN), mild cognitive impairment (MCI), or Alzheimer’s disease (AD) (note that some participants progressed from MCI to AD while being followed, with the last assessment used for case designation in the replication analysis, while they may be designated as beginning at the MCI stage in the following analysis). For clinical rate of progression analysis we used the Clinical Dementia Scale Sum of Boxes Score (CDRSB)^43^, a broad measure of clinical progression and impairment well-validated in multiple studies^44; 45^.

We used linear mixed-effects modeling to test whether variation in *TET2* predicts longitudinal clinical progression using R version 3.5.2. We covaried for baseline age, sex, education, and CDRSB score as well as *APOE ε*4 dose. The model was implemented as follows: Δ CDRSB = β_0_ + β_1_Age_baseline_ × Δt + β_2_Sex_female_ × Δt + β_3_Education_baseline_ × Δt + β_4_CDRSB_baseline_ × Δt + β_5_APOE 4_dose_ × Δt + β_6_*TET2*_carrier status_ × Δt + (1|subject) + ε

## RESULTS

Of the 1,539 samples in the original set, a total of 73 samples were removed from analysis after quality control. Two failed sex checks; 27 were pruned for relatedness; 12 were pruned due to an identifiable Mendelian variant (all of which were Sanger validated) meeting American College of Medical Genetics pathogenic or likely pathogenic criteria, including one *C9orf72* expansion carrier from the UAB set; one control was pruned for conversion to Mild Cognitive Impairment (MCI) after enrollment; and 31 cases were pruned because of phenotypic uncertainty or diagnosis of MCI or Parkinson’s disease (PD) rather than EOAD or FTD on re-evaluation after enrollment. The remaining dataset consisted of 1,466 individuals: 638 cases (294 EOAD and 344 FTD) and 828 controls. Of these cases and controls, 302 were of non-European ancestry (determined by principal component and admixture analysis, **Supplemental Figure 1**). Non-European ancestry individuals were excluded from the discovery set to reduce heterogeneity but were retained for replication. The resultant discovery set consisted of 1,164 individuals of European ancestry: 493 cases (228 EOAD and 265 FTD) and 671 controls. All demographic available information for each sample (case category, primary clinical diagnosis, sex, age at enrollment, *APOE* ε4 status, self-reported race/ethnicity, and principal components 1–4 and 5 ADMIXTURE coefficients) is available in **Supplemental Table 3**. The majority of cases were clinically diagnosed and did not have autopsy material available for neuropathological sub-grouping at the time of analysis. Primary clinical diagnoses included as AD were logopenic variant primary progressive aphasia (29), posterior cortical atrophy (26), frontal AD (17), language AD (17), vascular AD (8), AD + dementia with Lewy bodies (DLB) (5), and AD not otherwise specified (126). Primary clinical diagnoses included as FTD were behavioral variant FTD (83), corticobasal syndrome (65), nonfluent variant primary progressive aphasia (43), FTD + ALS (20), primary supranuclear palsy (17), semantic variant primary progressive aphasia (17), argyrophilic grain disease (5), and FTD not otherwise specified (15). The expected enrichment in *APOE* ε4 was observed in AD cases (58% with at least one *APOE* ε4 allele versus 28% in controls, \**p*=1.8×10^−16^ by Fisher’s exact test), but not in FTD (28% in FTD and controls, *p*=0.87).

We compared EOAD vs. control, FTD vs. control, or a combined analysis of EOAD and FTD vs. control across all variant filtering conditions (see Methods). In the discovery analysis of combined burden across EOAD and FTD vs. control, with variants absent from population databases and with a CADD score > 10 (including non-coding variants in GenoSkyline-Plus regions), one gene-disease association passed the multiple comparison significance threshold: *TET2* (SKAT uncorrected *p*=6.5×10^−8^, corrected *p*=0.0037; **Table 1**, model corrects for number of *APOE* ε4 alleles, sex, principal components 1–4, and 5 ADMIXTURE ancestral proportions). Note that, while we applied a multiple correction cutoff of 57,354 based on three main clusters of correlated filter conditions (**Supplemental Figure 2**), the *p* value for *TET2* would also pass a strict Bonferroni correction for 516,186 implicit tests (19,118 genes, 27 filter conditions) if we conservatively did not consider the correlated nature of the different filter sets (Bonferroni *p*=0.033). Statistical tests separately comparing EOAD vs. control and FTD vs. control did not pass the same degree of multiple testing correction but results for those comparisons are provided in **Supplemental Tables 4** (EOAD) **and 5** (FTD) and demonstrate that the nominal enrichment level in *TET2* is similar in both EOAD and FTD. No other gene reached even nominal significance (*p* < 1×10^−5^) in any filter condition, so *TET2* was the only gene considered for replication analysis. However, in the interest of making data from this study highly available, counts and *p* values for all genes assessed are provided in **Supplemental Table 6.**

**Table 1:** Discovery and replication for private, CADD > 10 coding and non-coding variants in *TET2* (combined analysis of all cases, AD and FTD, vs. controls). Variants in *TET2* absent from population databases and with a CADD score > 10 (including non-coding variants in GenoSkyline-Plus regions) in the combined analysis considering both EOAD and FTD cases vs. controls was the only qualifying gene and filter set in the discovery analysis to reach statistical significance. While we applied a correction factor of 57,354 based on genome wide (19,118 HGNC protein-coding genes) testing of three clusters of correlated filter conditions (**Supplemental Figure 2**), *TET2* remains significant if we conservatively do not consider the correlated nature of the different filter sets and instead apply a strict Bonferroni correction (*p* = 0.033). The primary test was SKAT corrected for number of *APOE* ε4 alleles, sex, principal components 1–4, and 5 ADMIXTURE ancestral proportions. Fisher’s exact test yielded similar raw *p* values and was highly correlated with SKAT (Pearson’s r=0.76 of log-transformed *p* values) and is presented here for consistency with Table 2, where some cohorts did not have covariate data available for SKAT and therefore relied on Fisher’s exact. The main analyses based on pre-determined criteria are bolded and * indicates significance (*p*<0.05 after correction). NA = Not Applicable. Replication cohorts are listed individually for reference as well as combined discovery plus replication statistics. Subsets of AD only vs. control and FTD only vs. control are provided in **Supplemental Table 4** and **Supplemental Table 5**, respectively.

| Cohort Type | Cohort | Cases w/ | Cases w/o | Ctrl s w/ | Ctrl s w/o | SKAT *p* | Corr. *p* | FET *p* | OR (95% CI) |
| --- | --- | --- | --- | --- | --- | --- | --- | --- | --- |
| Discovery | UCSF Eur. (EOAD & FTD) | 19 | 474 | 1 | 670 | 6.5x10<sup>-8</sup> | 3.7x10<sup>-3*</sup> | 8.1x10<sup>-7</sup> | 26.8 (4.2–1112) |
| Replication | UCSF Non-Eur. (EOAD & FTD) | 7 | 138 | 1 | 156 | 0.840 | NA | 0.031 | 7.9 (1.0–358) |
| Replication | ADSP Genomes (LOAD) | 64 | 2,144 | 36 | 2,172 | 4.6x10<sup>-4</sup> | NA | 6.0x10<sup>-3</sup> | 1.8 (1.2–2.8) |
| Replication | AMP-AD (LOAD & FTD) | 15 | 918 | 7 | 439 | 0.907 | NA | 1.000 | 1.0 (0.4–3.0) |
| Replication | All Replication Cohorts | 86 | 3,200 | 44 | 2,767 | 7.1x10<sup>-3</sup> | 7.1x10<sup>-3*</sup> | 4.4x10<sup>-3</sup> | 1.7 (1.2–2.5) |
| Combined | Discovery + All Replication | 105 | 3,674 | 45 | 3,437 | 8.0x10<sup>-6</sup> | NA | 7.0x10<sup>-6</sup> | 2.2 (1.5–3.2) |

All qualifying variants in cases for *TET2* were both Sanger validated and visually evaluated in the Integrative Genomics Viewer (IGV). Two variants failed Sanger validation (adjacent erroneous indel calls in a single sample) and were excluded from the variant counts in **Table 1**, all statistics, and in **Supplemental Table 7** where all qualifying variants in *TET2* are listed. In addition, two cases had adjacent variant calls that were found to make up one variant. These were also corrected in all statistical analyses and tables. The single control with a qualifying *TET2* variant did not have material available for Sanger sequencing but appeared valid in IGV (a single nucleotide variant with 8 alternate allele reads among 18 total reads).

Next, we assessed potential confounding due to stratification by a QQ plot of the *p*-value distribution for the filter set that produced the top result and observed no genomic inflation consistent with a well-matched case-control dataset (λ = 0.95, **Figure 1**).

To help inform the types of sequencing datasets to target for replication, we assessed the variant type (coding or non-coding) and associated disease for all qualifying *TET2* variants in the discovery set. Qualifying variants were observed in 11 EOAD cases, eight FTD cases, and one control. Of the 11 EOAD cases, one had depressive symptoms, one had language symptoms and possible corticobasal syndrome, one had logopenic variant primary progressive aphasia, and one had a previous diagnosis of behavioral variant FTD revised to frontal AD (seven had no additional noted phenotypes). Of the eight FTD cases, three had corticobasal syndrome (one of whom had AD symptoms and possible posterior cortical atrophy), one had FTD + ALS, and four had behavioral variant FTD (one with AD symptoms and one with seizures). Nine cases in total harbored coding variants, seven of which were canonical loss-of-function variants (four EOAD and three FTD). Because non-coding variants make up a large portion of the signal, we assessed coding and non-coding variants separately. We observed a similar level of enrichment for both coding and non-coding variants in EOAD and FTD cases when these types of variants were considered independently of one another (**Figure 2A**). Furthermore, the non-coding variants were prevalent in regions predicted to have regulatory function (**Figure 2B**). Combined with the high number of canonical loss-of-function variants, these data support a model whereby *TET2* haploinsufficiency, resulting from either canonical loss-of-function variation or expression-altering non-coding variation, may contribute risk to neurodegenerative disease.

To replicate this finding, we used six additional cohorts (five independent, one internal) with available sequencing data from patients diagnosed with a neurodegenerative disorder and healthy controls. Based on the variants discovered in *TET2*, we attempted to replicate the association between aggregate rare variant burden in *TET2* and disease risk using two arms: the same conditions used in discovery applied to other genome sequencing datasets as a primary measure, and canonical loss-of-function only analysis as a secondary measure to allow for incorporation of exome sequencing datasets. We assessed three cohorts with genome sequencing data for replication using the same conditions applied in the discovery set: ADSP (the Alzheimer’s Disease Sequencing Project) (2,208 late-onset AD (LOAD) cases and 2,208 controls), the Accelerating Medicines Partnership – Alzheimer’s Disease (AMP-AD) cohort (749 LOAD, 184 FTD, and 446 controls), and the non-European ancestry individuals from our cohort not assessed in the discovery set (66 EOAD, 79 FTD, and 157 controls). Assessment of these three cohorts revealed replication of the signal for *TET2* overall for early- and late-onset AD and FTD combined vs. control (*p*=0.0071; **Table 1**). Although the statistics for separate analyses of EOAD vs. control and FTD vs. control did not meet significance criteria, secondary analysis of those subgroups revealed similar levels of enrichment within each distinct condition (**Supplemental Table 4** (EOAD) and **Supplemental Table 5** (FTD)). Because of the established genetic overlap between FTD and ALS^46^, we also assessed variants in Project MinE^47^ (4,366 ALS cases and 1,832 controls) and observed a non-significant trend towards a slight enrichment in ALS cases (OR 1.3, 95% CI 1.1–2.7; **Supplemental Table 5**). While not a formal replication because no ALS cases were included in the discovery set, we present these findings in **Supplemental Table 5** along with FTD statistics.

Finally, we assessed predicted loss-of-function variants alone in all aforementioned cohorts (UCSF European discovery set, UCSF non-European replication set, ADSP, AMP-AD, Project MinE) along with exome sequencing data from a second ALS dataset^8^ and additional exome samples from ADSP^48^ for a total of seven sample sets. We observed a robust signal for association between predicted canonical loss-of-function variants and disease across multiple disease cohorts (**Table 2**). Specifically, three of the four largest independent replication cohorts (ADSP genomes (LOAD), ADSP exomes (LOAD), and HA-Duke-Stanford ALS exomes) all exhibit independent nominal replication (*p*<0.05). Meta-analysis of all canonical loss-of-function variants from all available cohorts (across EOAD, LOAD, FTD, and ALS) yields a *p*-value below commonly used exome-wide significance cutoffs (*p*=2.5×10^−7^; **Table 2**), and subgroup analyses of both AD and FTD–ALS vs. controls were each nominally significant (*p*<0.05) and suggest similar degrees of enrichment.

**Table 2:** Canonical loss-of-function variation in *TET2* is nominally enriched in both AD and FTD-ALS. Because of the high number of canonical loss-of-function variants in *TET2* observed in the discovery analysis, we performed a separate assessment of loss-of-function variants alone. Although the loss-of-function model did not pass multiple testing correction in the discovery analysis because of the low number of qualifying counts, *TET2* was the highest ranked loss-of-function gene (lowest *p*-value for enrichment in cases). Note the additional inclusion of ALS exomes (HA-Duke-Stanford). * indicates that the combined analysis across all cases and controls (in bold) was below an arbitrary exome-wide cutoff of 2.5×10^−6^ (a commonly used threshold based on correction of *p*<0.05 for ∼20,000 genes). SKAT values could not be calculated for ALS sets (and thus not for summed replication and discovery+replication sets) because necessary covariate data were not available for these cohorts, although both ALS cohorts were independently filtered to only individuals of European ancestry. Below this combined analysis, we also present summaries by each disease, which achieved nominal significance (*p* < 0.05) for both combined analysis of all AD cases and of all FTD and ALS cases. Note the addition of summary statistics from ADSP exomes in this section as well. For ADSP exomes, the *p*-value from the VEP HIGH meta-analysis model is shown (publicly available from NIAGADS). For comparison, we have also listed the frequency of *TET2* loss-of-function carriers in population databases (gnomAD minus TOPMed set added to counts from TOPMed), which is similar to the frequencies observed in control groups we analyzed. All frequencies are the percentage of individuals harboring a loss-of-function variant (not minor allele frequency) except ADSP exomes for which cumulative minor allele frequency (CMAF) for both cases and controls is listed.

| Cohort Type | Cohort | Cases w/ | Cases w/o | Ctrl's w/ | Ctrl's w/o | SKAT *p* | FET *p* | OR (95% CI) | Cases Freq. | Ctrl's Freq. |
| --- | --- | --- | --- | --- | --- | --- | --- | --- | --- | --- |
| Discovery | UCSF Eur. (EOAD & FTD) | 7 | 486 | 0 | 671 | 4.0x10<sup>-3</sup> | 2.4x10<sup>-3</sup> | ∞ (2–∞) | 1.42% | 0.00% |
| Replication | UCSF Non-Eur. (EOAD & FTD) | 1 | 144 | 1 | 156 | 0.058 | 1.000 | 1.1 (0.0–85.5) | 0.69% | 0.64% |
| Replication | ADSP Genomes (LOAD) | 31 | 2,177 | 12 | 2,196 | 3.7x10<sup>-4</sup> | 5.2x10<sup>-3</sup> | 2.6 (1.3–5.6) | 1.40% | 0.54% |
| Replication | AMP-AD (LOAD & FTD) | 0 | 933 | 1 | 445 | 0.215 | 0.323 | 0.0 (0.0–18.6) | 0.00% | 0.22% |
| Replication | Project MinE (ALS) | 21 | 4,345 | 5 | 1,827 | NA | 0.289 | 1.8 (0.6–6) | 0.48% | 0.27% |
| Replication | HA-Duke-Stanford (ALS) | 11 | 2,863 | 5 | 6,400 | NA | 2.0x10<sup>-3</sup> | 4.9 (1.6–18.1) | 0.38% | 0.08% |
| Replication | All Rep. Cohorts (AD, FTD, ALS) | 64 | 10,462 | 24 | 11,024 | NA | 8.1x10<sup>-6</sup> | 2.8 (1.7–4.7) | 0.61% | 0.22% |
| Combined | Discovery + Replication | 71 | 10,948 | 24 | 11,695 | NA | 2.5x10<sup>-7*</sup> | 3.2 (2.0–5.3) | 0.64% | 0.20% |
| Combined Subset | AD except ADSP Exomes | 35 | 3,216 | 14 | 3,468 | 2.7x10<sup>-3</sup> | 1.4x10<sup>-3</sup> | 2.7 (1.4–5.4) | 1.08% | 0.40% |
| Summary Stat. Set | ADSP Exomes (summary stats) | 6,345 total cases |  | 4,893 total controls |  | 0.019 (ADSP model *p* value) |  |  | CMAF 0.49% |  |
| Combined Subset | All FTD | 4 | 524 | 2 | 1,272 | 0.921 | 0.065 | 4.9 (0.7–53.7) | 0.76% | 0.16% |
| Combined Subset | All ALS | 32 | 7,208 | 10 | 8,227 | NA | 1.4x10<sup>-4</sup> | 3.7 (1.8–8.3) | 0.44% | 0.12% |
| Combined Subset | All FTD & ALS | 36 | 7,732 | 12 | 9,499 | NA | 3.0x10<sup>-5</sup> | 3.7 (1.9–7.8) | 0.46% | 0.13% |
| Population DBs | gnomAD+TOPMed | – | – | 284 | 196,035 | – | – | – | – | 0.14% |

To assess potential clinical implications of rare variation in *TET2*, we queried the Alzheimer’s Disease Neuroimaging Initiative (ADNI) dataset^40–42^, which includes clinical rate of progression data. We used linear mixed-effects modeling to test whether qualifying rare variation (based on the discovery condition that passed multiple corrections testing) in *TET2* predicts longitudinal clinical progression. We covaried for baseline age, sex, education, and CDRSB score as well as *APOE ε*4 dose. A total of 786 ADNI participants had *TET2* genotypes available for analysis. There was no significant difference in the distribution of *TET2* rare variant carriers by sex, *APOE ε*4 dose, education, or baseline CDRSB score (**Supplemental Table 8**). There was a significant difference between *TET2* rare variant carriers and non-carriers by baseline age (**Supplemental Table 8**) but recall that baseline age is corrected for as a covariate along with sex, education, *APOE ε*4 dose, and baseline CDRSB score. Using linear mixed effects regression, we found a significant relationship between carrying any *TET2* rare variant and clinical progression as measured by change in CDRSB score (β ± SE = 0.14 ± 0.06; \**p*=0.03) (**Figure 3**). A similar finding was observed when our analyses were limited to *TET2* loss-of-function variant carriers (β ± SE = 0.17 ± 0.09; \**p*=0.04) (**Supplemental Figure 3**) (although we corrected for covariates for rigor, no covariates were significantly associated with *TET2* loss-of-function carrier status (**Supplemental Table 9**)). We also explored whether rare variation in *TET2* predicted changes in CDRSB and cognition (measured by Mini Mental State Exam (MMSE) score changes^49^) in MCI and control when analyzed separately. To maximize sample size, we limited this analysis to all *TET2* variant carriers without further subdivision to loss-of-function variant carriers. When constraining the analysis to MCI, *TET2* rare variant carriers (n=8) demonstrated a greater CDRSB change over time compared to noncarriers of a higher magnitude and significance compared to the pooled analysis of control, MCI and AD (β ± SE = 0.64 ± 0.12; \**p*=6.17×10^−8^) when correcting for the covariates outlined above. Of note, *TET2* rare variant carriers diagnosed with MCI also demonstrated greater decreases in MMSE changes when compared to non-carriers (β ± SE = −0.47 ± 0.17; \**p*=0.01). Within controls (n=6), there were no significant associations between *TET2* variant carrier status and either CDRSB or MMSE.

**Figure 3:**
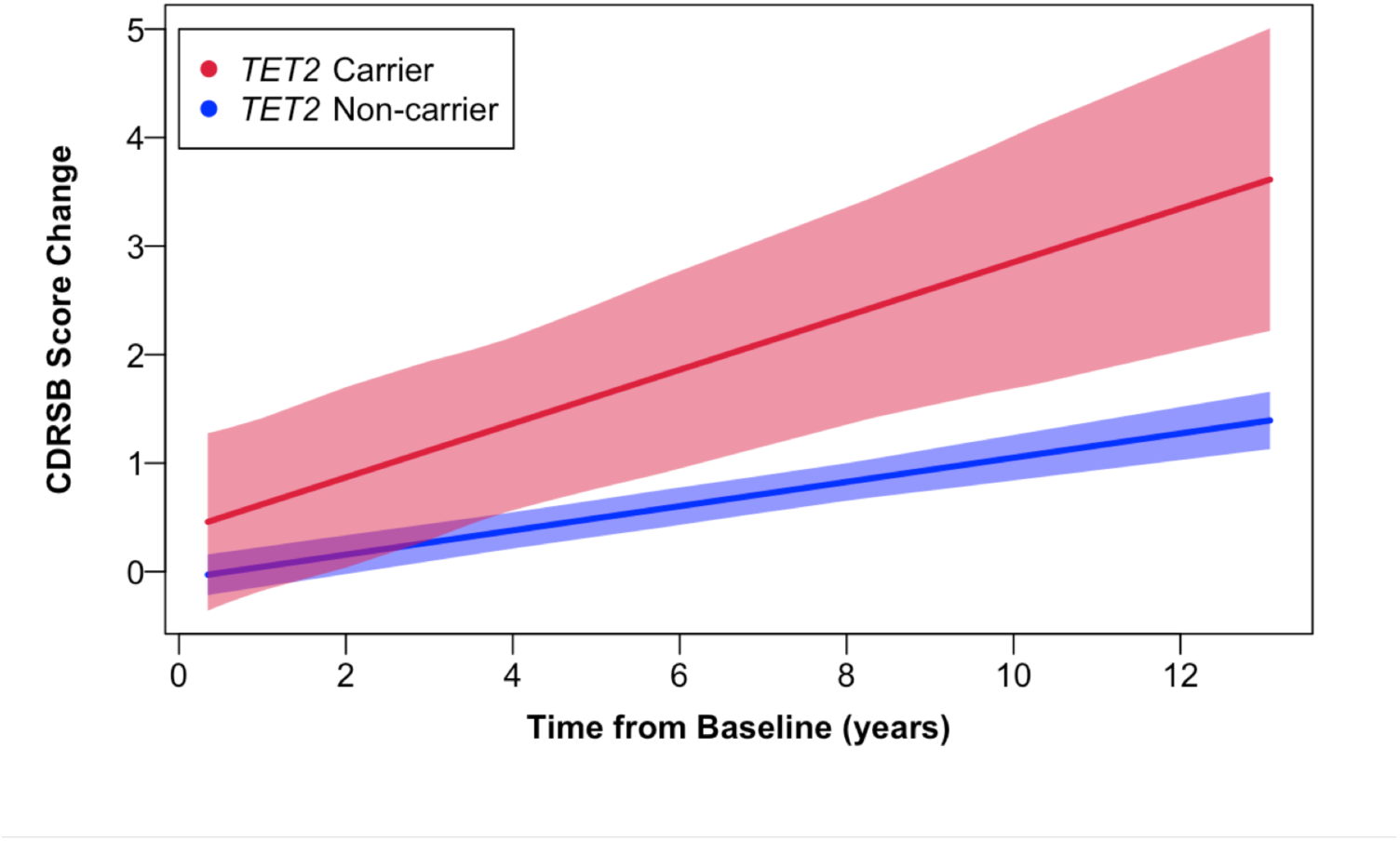
Longitudinal CDRSB changes in ADNI participants with qualifying *TET2* rare variants. *TET2* rare variant carriers show greater CDRSB changes over time compared to non-carriers after controlling for age, sex, education, *APOE ε*4, and baseline CDRSB score (β ± SE = 0.14 ± 0.06; \**p*=0.03). The lines depicted illustrate CDRSB change with 95% confidence intervals in shading. ADNI – Alzheimer’s Disease Neuroimaging Initiative; CDRSB – Clinical Dementia Rating Sum of Boxes Score.

## DISCUSSION

In this study, we identified a significant excess of rare, likely deleterious variation in *TET2* as a risk factor for multiple neurodegenerative disorders, including EOAD, LOAD, FTD, and ALS. This finding is important for two main reasons. First, *TET2* plays an important role in the conversion of methylation to 5-hydroxymethylation, implicating dysfunction in a pathway known to be critical during aging^50^ and learning and memory^51^ in age-associated diseases like LOAD and FTD. Second, it is striking that the effect sizes in both coding and non-coding variant enrichments were comparable. This point suggests that further investigation of non-coding variation in complex disease genome sequencing studies holds potential for the identification of new contributors to disease.

TET2 promotes de-methylation of DNA by catalyzing conversion of methylation to 5-hydroxymethylation, and is highly expressed in brain (reviewed in^52^). Defined methylation changes occur with age in humans (“Horvath’s clock”, reviewed in^50^) and there is some evidence for an association between increased “methylation age” and disease (systematically reviewed in ^53^). Taken together, this raises speculation that reduced function or loss of TET2, a critical regulator in methylation processes, may have adverse age-associated effects. Evidence from mouse models further supports this idea. Specifically, promoting the conversion of methylation to 5-hydroxymethylation by either exercise-induced upregulation of *TET2*^54^ or artificial overexpression of *TET2*^55^ improves memory in mice by increasing neurogenesis in the dentate gyrus. Conversely, reducing *TET2* in mouse hippocampus leads to reduced neurogenesis and impaired memory^55^, consistent with its role in promoting adult neurogenesis in mice^56^. Finally, reduction of *TET2* in mouse primary neurons also reduced cell survival^57^. All of these observations are consistent with detrimental consequences of loss of *TET2* function and suggest that neurons may be particularly vulnerable to these effects. Further support for a generally important role of TET enzymes comes from a preprint implicating mono- and bi-allelic loss-of-function of *TET3* in childhood diseases^58^ (*TET3* is more constrained against loss-of-function based on population database estimates^35^, which (along with bi-allelic contributions) could explain the earlier ages observed). In addition to general evidence for the importance of TET2 and other TET enzymes, an intriguing and more specific role for *TET2* has also been proposed in a preprint implicating *TET2* in microglial response, particularly around amyloid plaques^59^, suggesting that *TET2* loss-of-function may prevent its recruitment into a protective role (similar to recent findings on *TREM2* suggesting that higher secreted TREM2 levels are protective^60^, supporting a model where risk-conferring *TREM2* variants result in loss-of-function). Finally, the human data we analyzed from ADNI is consistent with deleterious consequences of *TET2* rare variants, with our observations supporting a faster rate of both general clinical decline (CDRSB) and cognitive decline (MMSE).

The strongest association signal in the discovery cohort was a combined analysis across all EOAD and FTD cases together. While we recognize the drawbacks of a combined analysis across phenotypes, we argue that the benefits outweigh the drawbacks for two critical reasons beyond the increase in sample size: (1) known effects of genetic pleiotropy, and (2) the possibility of identifying shared pathways between diseases.

The first reason supporting comparison across EOAD and FTD is that genetic pleiotropy—where a single locus contributes variance to multiple, different phenotypes—may play a role in neurodegenerative disease risk. Our group and others have provided support for this idea through several studies investigating multiple neurodegenerative diseases using GWAS approaches^11–15^. In addition to common risk variants, there is clear evidence of moderately to highly penetrant rare variation in single genes conferring risk or causality for multiple neurodegenerative diseases. First, rare variants in *TREM2* confer risk for both AD^61^ and FTD^62^. Second, rare variation in multiple established genes such as *TBK1* and *C9orf72* confer risk or are causative across the ALS-FTD spectrum^63^. Third, moderately penetrant common risk alleles like *APOE* ε4 are primarily associated with AD, but also associated with risk for Dementia with Lewy Bodies (DLB)^64^, FTD^12^, and age of onset in *C9orf72* carriers^65^. Fourth, *GBA* and *SNCA* were first identified as risk factors for Parkinson’s disease (PD), but also confer risk for DLB^64^. Finally, rare pathogenic variants in *MAPT* typically cause FTD^66; 67^, but the R406W pathogenic variant has also been associated with an EOAD presentation^68; 69^. Furthermore, common variants near *MAPT* (tagging the H1 haplotype, associated with higher tau expression^70^) are associated with AD, PD, FTD, and ALS^11; 12; 15^.

A second important reason to analyze across different patient populations is that performing analyses across cohorts of patients diagnosed with different neurodegenerative disorders but with partially overlapping underlying neuropathology (i.e., tau-containing protein aggregates in both AD and approximately half of FTD cases; TDP-43 containing protein aggregates in ALS, approximately half of FTD cases, and some AD cases) may identify shared dysregulated pathways, and has the potential to identify therapeutic targets with relevance across multiple neurodegenerative diseases. Indeed, our discovery that rare variation in *TET2* is associated with multiple neurodegenerative diseases suggests age-related changes in methylation may be relevant across a broad spectrum of neurodegeneration.

In conclusion, we provide evidence that loss of *TET2* function confers risk for EOAD, LOAD, FTD, and ALS. Specifically, we found that, in aggregate, both coding and non-coding qualifying rare variation in *TET2* is associated with approximately a 2-fold risk increase across diverse populations of patients with AD, FTD, and ALS, and that canonical loss-of-function variation in *TET2* is associated with approximately a 3-fold risk increase for these diseases. We note that, similar to any burden test, it is impossible from aggregate enrichment values to deconvolute variable penetrance levels among disease-relevant alleles and the degree of enrichment for truly associated variation. Future work assessing the functional effects of particular alleles and their concomitant levels of risk to individual variant carriers would be helpful in this regard. Additionally, further work is required to understand the local and global mechanisms by which alterations to TET2 levels and/or function contributes to disease risk, whether this risk is anchored to TET2’s effects on aging biology, and, if so, whether rare variation in *TET2* also confers risk to other age-associated neurodegenerative diseases.

## SUPPLEMENTAL DATA DESCRIPTION

Supplemental data includes three figures and nine tables. The supplemental figures and two of the supplemental tables (4,5,8,9) are provided in the **Supplemental Data**. Four supplemental tables are provided in an Excel file (1,2,3,7), and 1 supplemental table (6) is provided as a zipped text file.

## Data availability

All data in both discovery and replication sets are available through either an application for access by qualified researchers, or through public availability. In addition to providing information on how to access all datasets below, we have also included a supplemental text file with count data for all conditions assessed in the discovery cohort (**Supplemental Table 6**). Genome data for UCSF-enrolled participants is available to qualified researchers on request to the UCSF Memory and Aging Center (https://memory.ucsf.edu/research-trials/professional/open-science). Genome data from University of Alabama at Birmingham–enrolled participants is available from the National Institute on Aging Genetics of Alzheimer’s Disease Data Storage (NIAGADS) site under project NG00082. Data from control subjects sequenced at HudsonAlpha are available under dbGaP accessions phs001625.v1.p1 and phs001089.v3.p1. ADNI (Alzheimer’s Disease Neuroimaging Initiative, part of the ADSP genomes batch call) and ADSP data are available at NIAGADS under projects NG00066 and NG00067 and on dbGaP under accession phs000572.v7.p4 (see Supplemental Extended Acknowledgements for all ADSP investigators and funding sources). Data from AMP-AD are available through controlled access to the AMP-AD Knowledge Portal on Synapse (DOI: 10.7303/syn2580853). Data from Project MinE are freely available online (http://databrowser.projectmine.com/^71^. Summary statistics from^8^ are freely available online. 1000 genomes data are freely available online (http://www.internationalgenome.org^27^.

## Authors’ contributions

Experimental design: JNC, EGG, LWB, GMC, BNL, GDR, BLM, RMM, JSY. Data collection: LWB, MDA, MLT, AMK, EDR, GDR, BLM, JSY. Data analysis and interpretation: JNC, EGG, LWB, JSN, BNL, GMC, RMM, JSY. Subject recruitment: EDR, GDR, BLM. Technical or administrative support: JSN, LWB, AMK. Writing the manuscript: JNC, EGG, LWB, JSY. Editing the manuscript: JNC, EGG, LWB, MLT, BNL, EDR, GDR, RMM, JSY.

## Acknowledgements

We thank the Genomic Services Lab at HudsonAlpha for DNA isolations, library generation, quality control and sequencing. We also acknowledge a key contribution from the availability of datasets through controlled access, without which replication for this study would not have been possible. In addition to referencing how to access these sets in the *Data availability* section, we reiterate our appreciation for these datasets here: ADSP, ADNI, the AMP-AD Knowledge Portal (doi:10.7303/syn2580853) (with contributions from ROSMAP^72^, Mayo^73^, and MSBB), the Project MinE Sequencing Consortium^47^, and HA-Duke-Stanford ALS exomes^8^. More detail on contributing studies is listed below, and the full list of contributors for ADSP and ADNI as listed in the Supplemental Acknowledgements because of space limitations.

CSER: dbGaP accession phs001089.v3.p1 contains data generated by the Clinical Sequencing Exploratory Research (CSER) Consortium established by the NHGRI. Funding support for CSER was provided through cooperative agreements with the NHGRI and NCI through grant numbers U01 HG007301 (Genomic Diagnosis in Children with Developmental Delay). Information about CSER and the investigators and institutions who comprise the CSER consortium can be found at https://cser-consortium.org.

NIMH: The results published here are in whole or part based upon data generated by the Pritzker Neuropsychiatric Disorders Research Consortium at the HudsonAlpha Institute for Biotechnology. Funding support was provided by the Pritzker Neuropsychiatric Disorders Research Consortium. A case-control study was conducted to assay rare, predicted deleterious genetic variants in patients who have psychotic major depression. The data that is provided is genome sequencing of the control samples (unaffected) from this study, which originated from the Molecular Genetics of Schizophrenia samples (Study #29 in the NIMH repository).

ROSMAP: Study data were provided by the Rush Alzheimer’s Disease Center, Rush University Medical Center, Chicago. Data collection was supported through funding by NIA grants P30AG10161, R01AG15819, R01AG17917, R01AG30146, R01AG36836, U01AG32984, U01AG46152, U01AG61356, the Illinois Department of Public Health, and the Translational Genomics Research Institute.

Mayo RNA-Seq genome sequencing data: Study data were provided by the following sources: The Mayo Clinic Alzheimers Disease Genetic Studies, led by Dr. Nilufer Taner and Dr. Steven G. Younkin, Mayo Clinic, Jacksonville, FL using samples from the Mayo Clinic Study of Aging, the Mayo Clinic Alzheimers Disease Research Center, and the Mayo Clinic Brain Bank. Data collection was supported through funding by NIA grants P50 AG016574, R01 AG032990, U01 AG046139, R01 AG018023, U01 AG006576, U01 AG006786, R01 AG025711, R01 AG017216, R01 AG003949, NINDS grant R01 NS080820, CurePSP Foundation, and support from Mayo Foundation. Study data includes samples collected through the Sun Health Research Institute Brain and Body Donation Program of Sun City, Arizona. The Brain and Body Donation Program is supported by the National Institute of Neurological Disorders and Stroke (U24 NS072026 National Brain and Tissue Resource for Parkinsons Disease and Related Disorders), the National Institute on Aging (P30 AG19610 Arizona Alzheimers Disease Core Center), the Arizona Department of Health Services (contract 211002, Arizona Alzheimers Research Center), the Arizona Biomedical Research Commission (contracts 4001, 0011, 05-901 and 1001 to the Arizona Parkinson’s Disease Consortium) and the Michael J. Fox Foundation for Parkinsons Research.

MSBB: These data were generated from postmortem brain tissue collected through the Mount Sinai VA Medical Center Brain Bank (MSBB) and were provided by Dr. Eric Schadt from Mount Sinai School of Medicine.

## SUPPLEMENTAL DATA

### SUPPLEMENTAL FIGURES

**Supplemental Figure 1:**
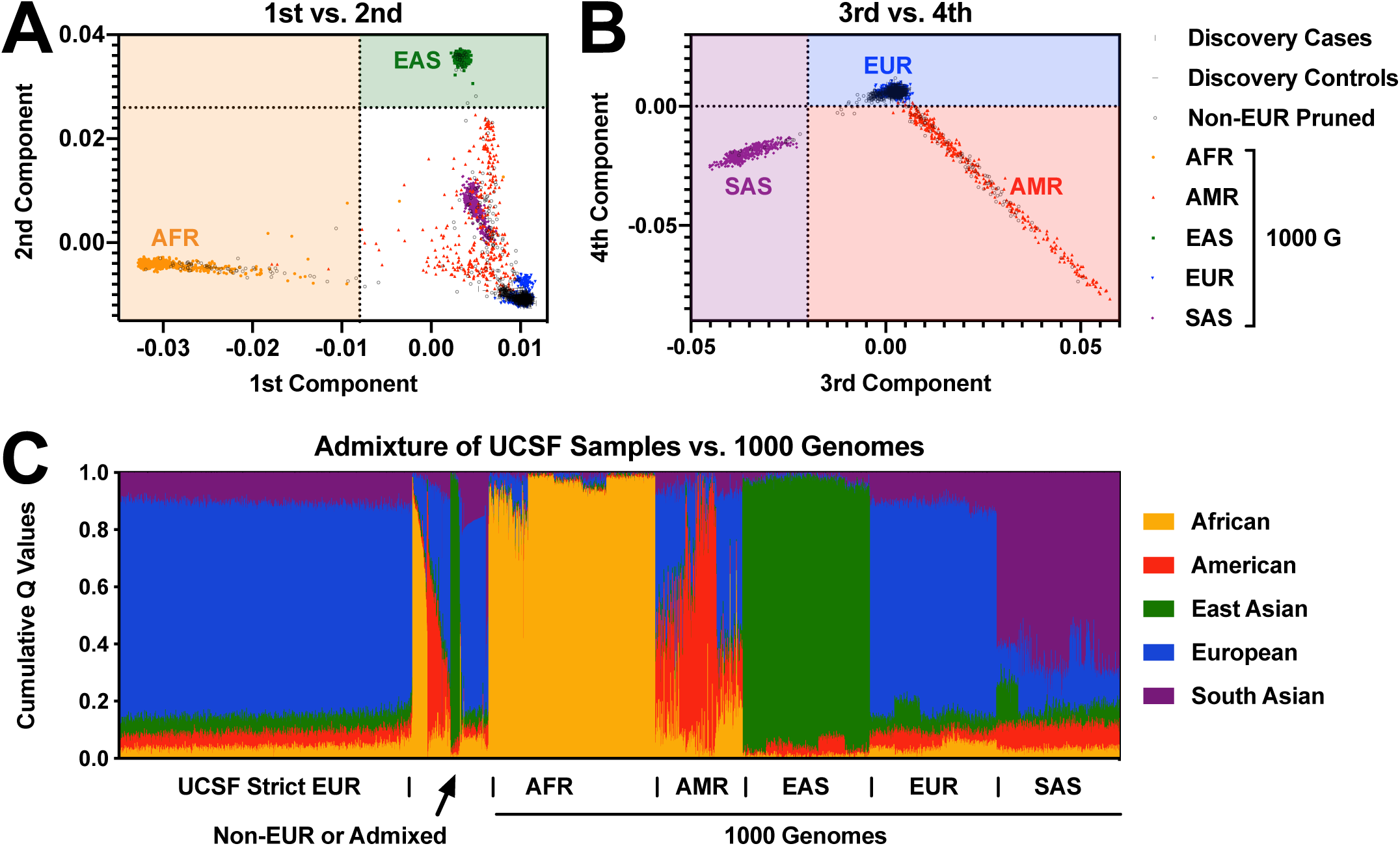
Ancestry analysis. Principal component analysis (PCA) of common variants was used to separate superpopulations by comparing clusters to 1000 genomes data for **A.** PC1 (separates AFR ancestry) and PC2 (separates EAS ancestry), and **B.** PC3 (separates SAS ancestry) and PC4 (separates AMR ancestry). **C.** The remaining samples were considered as candidates for EUR ancestry but were further pruned to disallow any ancestral proportion of greater than 15% by ADMIXTURE with k=5 from the 4 ancestral populations least enriched in EUR samples, again comparing to 1000 genomes data. The remaining samples are in the “UCSF Strict EUR” bin. Cases and controls are plotted as vertical and horizontal lines and cluster above the 1000 genomes control data in **B** and **C**, with samples pruned for non-EUR ancestry shown as circles.

**Supplemental Figure 2:**
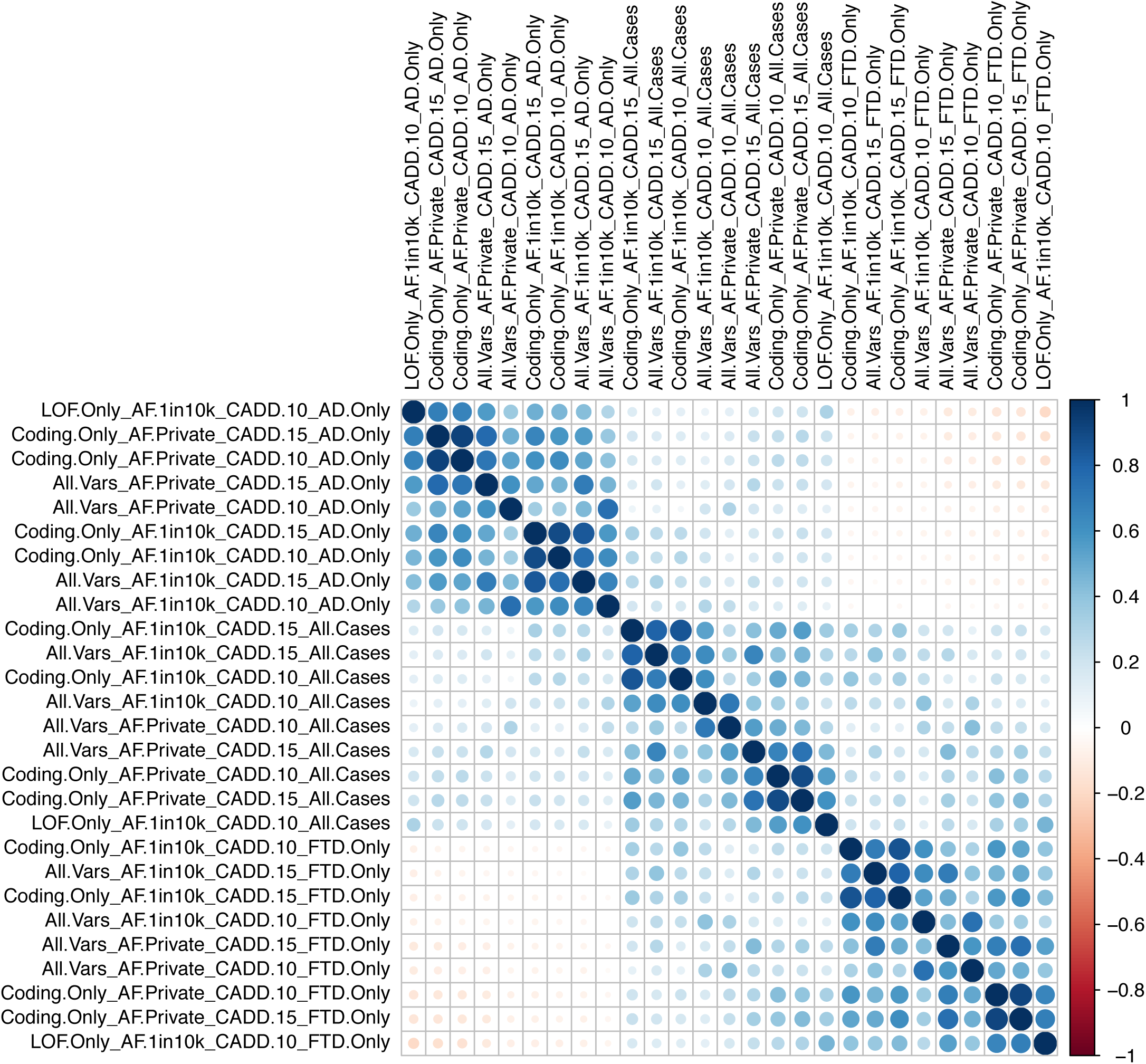
Cross-correlation plot of tested filter conditions. Pearson correlations were calculated for log transformed *p* values between all filter sets tested. Filter sets were positively correlated with one another within three main clusters emerging corresponding to case-control comparison groups.

**Supplemental Figure 3:**
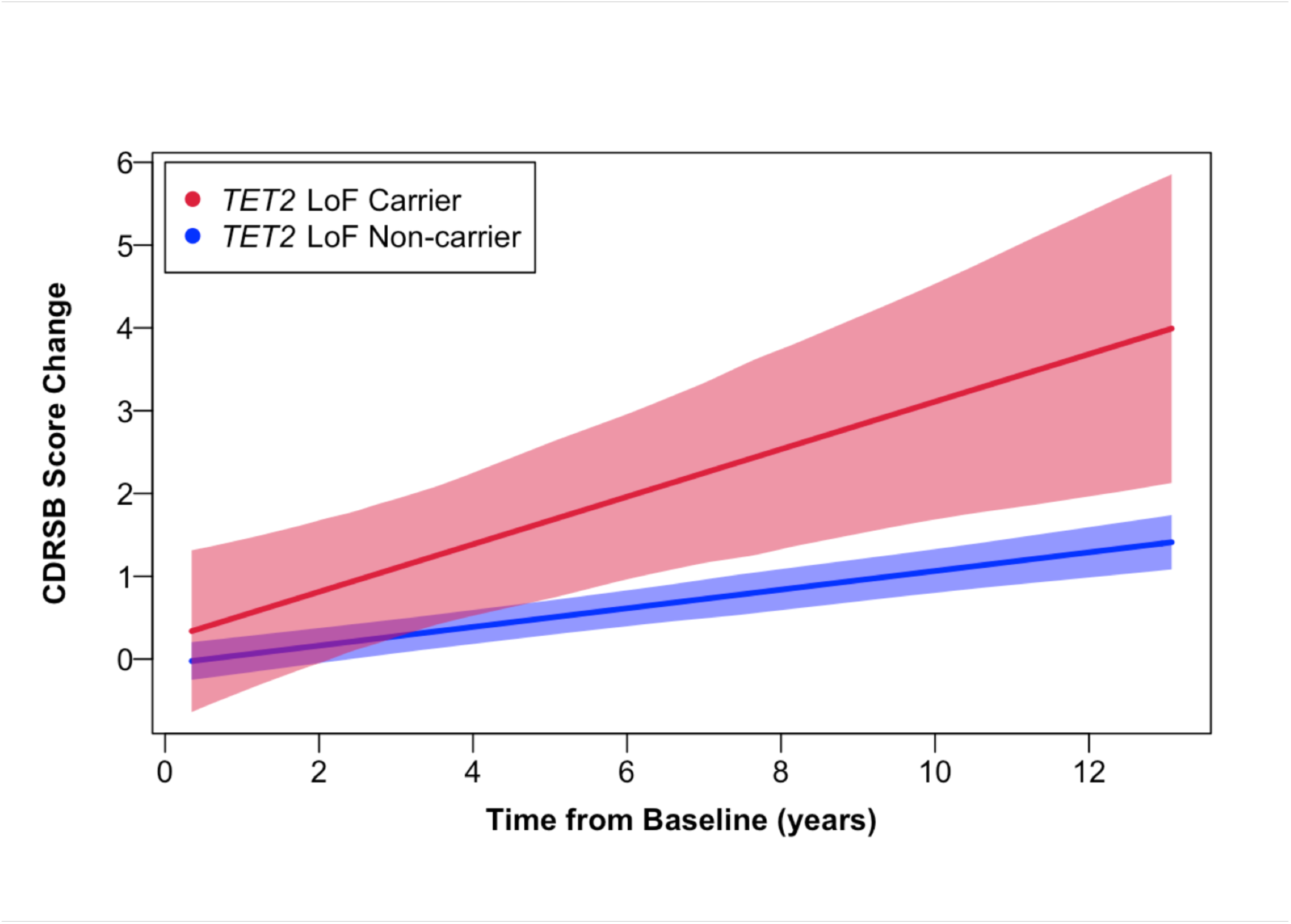
Longitudinal CDRSB changes in ADNI participants with *TET2* loss-of-function variants. *TET2* rare variant carriers whose changes are predicted to result in a loss-of-function variant show greater CDRSB changes over time compared to non-carriers after controlling for age, sex, education, *APOE ε*4, and baseline CDRSB score (β ± SE = 0.17 ± 0.09; \**p*=0.04). The lines depicted illustrate CDRSB change with 95% confidence intervals in shading. ADNI – Alzheimer’s Disease Neuroimaging Initiative; CDRSB – Clinical Dementia Rating Sum of Boxes Score.

### SUPPLEMENTAL TABLES

**Supplemental Table 1:** GenoSkyline-Plus tracks. Only tracks from tissue were included. See Excel file.

**Supplemental Table 2:** Demographic information. See Excel file.

**Supplemental Table 3:** List of all 19,118 hg19 HUGO Gene Nomenclature Committee (HGNC) genes tested. See Excel file.

**Supplemental Table 4:**
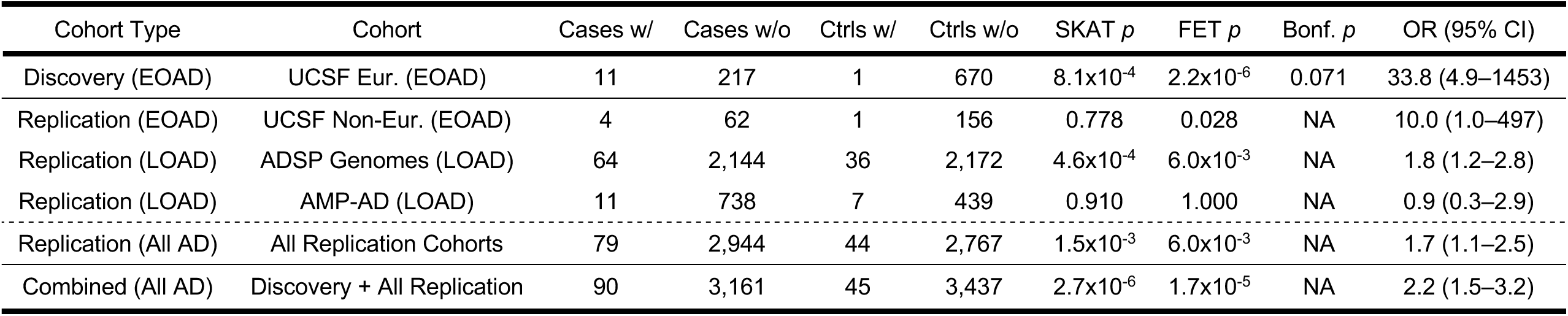
Discovery and replication for private, CADD > 10 coding and non-coding variants in *TET2* (analysis of AD vs. controls). Related to Table 1. Note that the discovery comparison did not meet the multiple corrections cutoff of *p*<0.05, therefore no formal replication was performed, so all statistics provided here are nominal.

| Cohort Type | Cohort | Cases w/ | Cases w/o | Ctrls w/ | Ctrls w/o | SKAT *p* | FET *p* | Bonf. *p* | OR (95% CI) |
| --- | --- | --- | --- | --- | --- | --- | --- | --- | --- |
| Discovery (EOAD) | UCSF Eur. (EOAD) | 11 | 217 | 1 | 670 | 8.1x10<sup>-4</sup> | 2.2x10<sup>-6</sup> | 0.071 | 33.8 (4.9–1453) |
| Replication (EOAD) | UCSF Non-Eur. (EOAD) | 4 | 62 | 1 | 156 | 0.778 | 0.028 | NA | 10.0 (1.0–497) |
| Replication (LOAD) | ADSP Genomes (LOAD) | 64 | 2,144 | 36 | 2,172 | 4.6x10<sup>-4</sup> | 6.0x10<sup>-3</sup> | NA | 1.8 (1.2–2.8) |
| Replication (LOAD) | AMP-AD (LOAD) | 11 | 738 | 7 | 439 | 0.910 | 1.000 | NA | 0.9 (0.3–2.9) |
| Replication (All AD) | All Replication Cohorts | 79 | 2,944 | 44 | 2,767 | 1.5x10<sup>-3</sup> | 6.0x10<sup>-3</sup> | NA | 1.7 (1.1–2.5) |
| Combined (All AD) | Discovery + All Replication | 90 | 3,161 | 45 | 3,437 | 2.7x10<sup>-6</sup> | 1.7x10<sup>-5</sup> | NA | 2.2 (1.5–3.2) |

**Supplemental Table 5:**
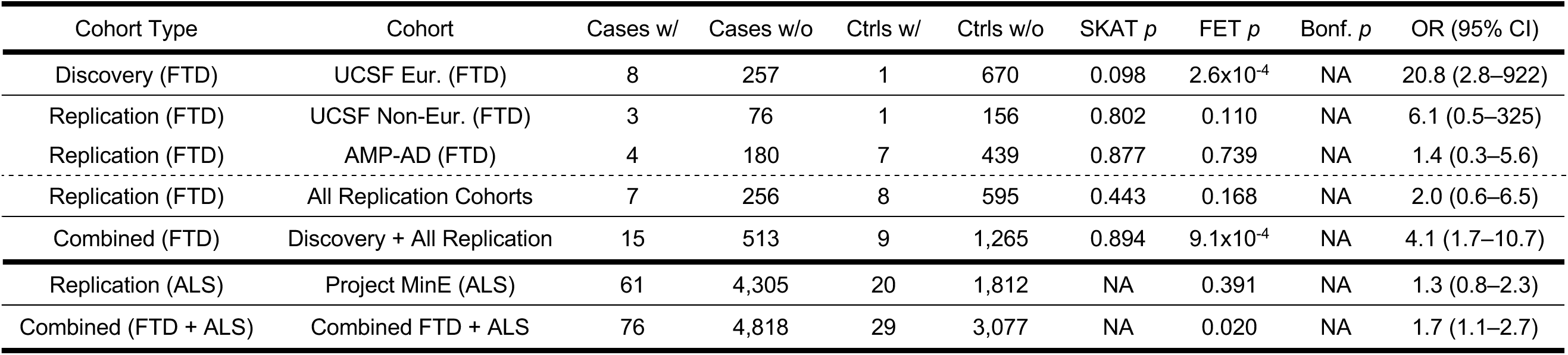
Discovery and replication for private, CADD > 10 coding and non-coding variants in *TET2* (analysis of FTD vs. controls). Related to Table 1. Note that the discovery comparison did not meet the multiple corrections cutoff of *p*<0.05, therefore no formal replication was performed, so all statistics provided here are nominal.

**Supplemental Table 6:** Case and control counts for all filter sets and genes for the discovery cohort. A header in this file describes how to obtain counts for any desired filter set and gene. See zipped text file.

**Supplemental Table 7:** Qualifying variants in *TET2* from all four cohorts with individual genotype data available. See Excel file.

**Supplemental Table 8:** ADNI Cohort Characteristics for Qualifying *TET2* Rare Variant Carriers.

|  | *TET2* Carrier Status |  | P-value |
| --- | --- | --- | --- |
|  | Non-Carrier | Carrier |  |
| N | 771 | 15 |  |
| Age (Years; Mean ± SD) | 73.2 ± 7.1 | 77.1 ± 6.5 | 0.03 |
| Sex (# Male (%)) | 433 (56.2%) | 7 (46.7%) | 0.64 |
| *APOE* ε4 Dose (%) |  |  | 0.81 |
| 0 | 451 (58.5%) | 10 (66.7%) |  |
| 1 | 266 (34.5%) | 4 (26.7%) |  |
| 2 | 54 (7.0%) | 1 (6.7%) |  |
| Diagnosis (#, %) |  |  | 0.89 |
| CN | 266 (34.5%) | 6 (40.0%) |  |
| MCI | 461 (59.8%) | 8 (53.3%) |  |
| AD | 44 (5.7%) | 1 (6.7%) |  |
| Education (Years; Mean ± SD) | 16.1 ± 2.8 | 16.3 ± 2.9 | 0.81 |
| CDRSB Baseline (Mean ± SD) | 1.1 ± 1.3 | 1.0 ± 1.2 | 0.64 |

CN—Normal Control; MCI – Mild Cognitive Impairment; AD – Alzheimer’s disease; SD – Standard Deviation; CDRSB – Clinical Dementia Rating Sum of Boxes Score.

**Supplemental Table 9:** ADNI Cohort Characteristics for *TET2* Loss-of-Function Carriers.

|  | *TET2* LoF Carrier Status |  | P-value |
| --- | --- | --- | --- |
|  | LoF Non-Carrier | LoF Carrier |  |
| N | 778 | 8 |  |
| Age (Years; Mean ± SD) | 73.2 ± 7.1 | 77.7 ± 5.7 | 0.08 |
| Sex (# Male (%)) | 436 (56.0%) | 4 (50.0%) | 1 |
| *APOE* ε4 Dose (%) | | | 0.57 |
| 0 | 455 (58.5%) | 6 (75.0%) |  |
| 1 | 268 (34.4%) | 2 (25.0%) |  |
| 2 | 55 (7.1%) | 0 (0.0%) |  |
| Diagnosis (#, %) |  |  | 0.61 |
| CN | 270 (34.7%) | 2 (25.0%) |  |
| MCI | 463 (59.5%) | 6 (75.0%) |  |
| AD | 45 (5.8%) | 0 (0.0%) |  |
| Education (Years; Mean ± SD) | 16.1 ± 2.8 | 16.5 ± 2.6 | 0.68 |
| CDRSB Baseline (Mean ± SD) | 1.1 ± 1.3 | 0.9 ± 0.7 | 0.59 |

CN—Normal Control; MCI – Mild Cognitive Impairment; AD – Alzheimer’s disease; SD – Standard Deviation; CDRSB – Clinical Dementia Rating Sum of Boxes Score.

## SUPPLEMENTAL ACKNOWLEDGEMENTS

### ADSP

The Alzheimer’s Disease Sequencing Project (ADSP) is comprised of two Alzheimer’s Disease (AD) genetics consortia and three National Human Genome Research Institute (NHGRI) funded Large Scale Sequencing and Analysis Centers (LSAC). The two AD genetics consortia are the Alzheimer’s Disease Genetics Consortium (ADGC) funded by NIA (U01 AG032984), and the Cohorts for Heart and Aging Research in Genomic Epidemiology (CHARGE) funded by NIA (R01 AG033193), the National Heart, Lung, and Blood Institute (NHLBI), other National Institute of Health (NIH) institutes and other foreign governmental and non-governmental organizations. The Discovery Phase analysis of sequence data is supported through UF1AG047133 (to Drs. Schellenberg, Farrer, Pericak-Vance, Mayeux, and Haines); U01AG049505 to Dr. Seshadri; U01AG049506 to Dr. Boerwinkle; U01AG049507 to Dr. Wijsman; and U01AG049508 to Dr. Goate and the Discovery Extension Phase analysis is supported through U01AG052411 to Dr. Goate, U01AG052410 to Dr. Pericak-Vance and U01 AG052409 to Drs. Seshadri and Fornage. Data generation and harmonization in the Follow-up Phases is supported by U54AG052427 (to Drs. Schellenberg and Wang).

The ADGC cohorts include: Adult Changes in Thought (ACT), the Alzheimer’s Disease Centers (ADC), the Chicago Health and Aging Project (CHAP), the Memory and Aging Project (MAP), Mayo Clinic (MAYO), Mayo Parkinson’s Disease controls, University of Miami, the Multi-Institutional Research in Alzheimer’s Genetic Epidemiology Study (MIRAGE), the National Cell Repository for Alzheimer’s Disease (NCRAD), the National Institute on Aging Late Onset Alzheimer’s Disease Family Study (NIA-LOAD), the Religious Orders Study (ROS), the Texas Alzheimer’s Research and Care Consortium (TARC), Vanderbilt University/Case Western Reserve University (VAN/CWRU), the Washington Heights-Inwood Columbia Aging Project (WHICAP) and the Washington University Sequencing Project (WUSP), the Columbia University Hispanic-Estudio Familiar de Influencia Genetica de Alzheimer (EFIGA), the University of Toronto (UT), and Genetic Differences (GD).

The CHARGE cohorts are supported in part by National Heart, Lung, and Blood Institute (NHLBI) infrastructure grant HL105756 (Psaty), RC2HL102419 (Boerwinkle) and the neurology working group is supported by the National Institute on Aging (NIA) R01 grant AG033193. The CHARGE cohorts participating in the ADSP include the following: Austrian Stroke Prevention Study (ASPS), ASPS-Family study, and the Prospective Dementia Registry-Austria (ASPS/PRODEM-Aus), the Atherosclerosis Risk in Communities (ARIC) Study, the Cardiovascular Health Study (CHS), the Erasmus Rucphen Family Study (ERF), the Framingham Heart Study (FHS), and the Rotterdam Study (RS). ASPS is funded by the Austrian Science Fond (FWF) grant number P20545-P05 and P13180 and the Medical University of Graz. The ASPS-Fam is funded by the Austrian Science Fund (FWF) project I904),the EU Joint Programme - Neurodegenerative Disease Research (JPND) in frame of the BRIDGET project (Austria, Ministry of Science) and the Medical University of Graz and the Steiermärkische Krankenanstalten Gesellschaft. PRODEM-Austria is supported by the Austrian Research Promotion agency (FFG) (Project No. 827462) and by the Austrian National Bank (Anniversary Fund, project 15435. ARIC research is carried out as a collaborative study supported by NHLBI contracts (HHSN268201100005C, HHSN268201100006C, HHSN268201100007C, HHSN268201100008C, HHSN268201100009C, HHSN268201100010C, HHSN268201100011C, and HHSN268201100012C). Neurocognitive data in ARIC is collected by U01 2U01HL096812, 2U01HL096814, 2U01HL096899, 2U01HL096902, 2U01HL096917 from the NIH (NHLBI, NINDS, NIA and NIDCD), and with previous brain MRI examinations funded by R01-HL70825 from the NHLBI. CHS research was supported by contracts HHSN268201200036C, HHSN268200800007C, N01HC55222, N01HC85079, N01HC85080, N01HC85081, N01HC85082, N01HC85083, N01HC85086, and grants U01HL080295 and U01HL130114 from the NHLBI with additional contribution from the National Institute of Neurological Disorders and Stroke (NINDS). Additional support was provided by R01AG023629, R01AG15928, and R01AG20098 from the NIA. FHS research is supported by NHLBI contracts N01-HC-25195 and HHSN268201500001I. This study was also supported by additional grants from the NIA (R01s AG054076, AG049607 and AG033040 and NINDS (R01 NS017950). The ERF study as a part of EUROSPAN (European Special Populations Research Network) was supported by European Commission FP6 STRP grant number 018947 (LSHG-CT-2006-01947) and also received funding from the European Community’s Seventh Framework Programme (FP7/2007-2013)/grant agreement HEALTH-F4-2007-201413 by the European Commission under the programme “Quality of Life and Management of the Living Resources” of 5th Framework Programme (no. QLG2-CT-2002-01254). High-throughput analysis of the ERF data was supported by a joint grant from the Netherlands Organization for Scientific Research and the Russian Foundation for Basic Research (NWO-RFBR 047.017.043). The Rotterdam Study is funded by Erasmus Medical Center and Erasmus University, Rotterdam, the Netherlands Organization for Health Research and Development (ZonMw), the Research Institute for Diseases in the Elderly (RIDE), the Ministry of Education, Culture and Science, the Ministry for Health, Welfare and Sports, the European Commission (DG XII), and the municipality of Rotterdam. Genetic data sets are also supported by the Netherlands Organization of Scientific Research NWO Investments (175.010.2005.011, 911-03-012), the Genetic Laboratory of the Department of Internal Medicine, Erasmus MC, the Research Institute for Diseases in the Elderly (014-93-015; RIDE2), and the Netherlands Genomics Initiative (NGI)/Netherlands Organization for Scientific Research (NWO) Netherlands Consortium for Healthy Aging (NCHA), project 050-060-810. All studies are grateful to their participants, faculty and staff. The content of these manuscripts is solely the responsibility of the authors and does not necessarily represent the official views of the National Institutes of Health or the U.S. Department of Health and Human Services.

The four LSACs are: the Human Genome Sequencing Center at the Baylor College of Medicine (U54 HG003273), the Broad Institute Genome Center (U54HG003067), The American Genome Center at the Uniformed Services University of the Health Sciences (U01AG057659), and the Washington University Genome Institute (U54HG003079).

Biological samples and associated phenotypic data used in primary data analyses were stored at Study Investigators institutions, and at the National Cell Repository for Alzheimer’s Disease (NCRAD, U24AG021886) at Indiana University funded by NIA. Associated Phenotypic Data used in primary and secondary data analyses were provided by Study Investigators, the NIA funded Alzheimer’s Disease Centers (ADCs), and the National Alzheimer’s Coordinating Center (NACC, U01AG016976) and the National Institute on Aging Genetics of Alzheimer’s Disease Data Storage Site (NIAGADS, U24AG041689) at the University of Pennsylvania, funded by NIA, and at the Database for Genotypes and Phenotypes (dbGaP) funded by NIH. This research was supported in part by the Intramural Research Program of the National Institutes of health, National Library of Medicine. Contributors to the Genetic Analysis Data included Study Investigators on projects that were individually funded by NIA, and other NIH institutes, and by private U.S. organizations, or foreign governmental or nongovernmental organizations.

### ADNI

Michael Weiner, MD (UC San Francisco, Principal Investigator, Executive Committee); Paul Aisen, MD (UC San Diego, ADCS PI and Director of Coordinating Center Clinical Core, Executive Committee, Clinical Core Leaders); Ronald Petersen, MD, PhD (Mayo Clinic, Rochester, Executive Committee, Clinical Core Leader); Clifford R. Jack, Jr., MD (Mayo Clinic, Rochester, Executive Committee, MRI Core Leader); William Jagust, MD (UC Berkeley, Executive Committee; PET Core Leader); John Q. Trojanowki, MD, PhD (U Pennsylvania, Executive Committee, Biomarkers Core Leader); Arthur W. Toga, PhD (USC, Executive Committee, Informatics Core Leader); Laurel Beckett, PhD (UC Davis, Executive Committee, Biostatistics Core Leader); Robert C. Green, MD, MPH (Brigham and Women’s Hospital, Harvard Medical School, Executive Committee and Chair of Data and Publication Committee); Andrew J. Saykin, PsyD (Indiana University, Executive Committee, Genetics Core Leader); John Morris, MD (Washington University St. Louis, Executive Committee, Neuropathology Core Leader); Leslie M. Shaw (University of Pennsylvania, Executive Committee, Biomarkers Core Leader); Enchi Liu, PhD (Janssen Alzheimer Immunotherapy, ADNI two Private Partner Scientific Board Chair); Tom Montine, MD, PhD (University of Washington); Ronald G. Thomas, PhD (UC San Diego); Michael Donohue, PhD (UC San Diego); Sarah Walter, MSc (UC San Diego); Devon Gessert (UC San Diego); Tamie Sather, MS (UC San Diego,); Gus Jiminez, MBS (UC San Diego); Danielle Harvey, PhD (UC Davis;); Michael Donohue, PhD (UC San Diego); Matthew Bernstein, PhD (Mayo Clinic, Rochester); Nick Fox, MD (University of London); Paul Thompson, PhD (USC School of Medicine); Norbert Schuff, PhD (UCSF MRI); Charles DeCArli, MD (UC Davis); Bret Borowski, RT (Mayo Clinic); Jeff Gunter, PhD (Mayo Clinic); Matt Senjem, MS (Mayo Clinic); Prashanthi Vemuri, PhD (Mayo Clinic); David Jones, MD (Mayo Clinic); Kejal Kantarci (Mayo Clinic); Chad Ward (Mayo Clinic); Robert A. Koeppe, PhD (University of Michigan, PET Core Leader); Norm Foster, MD (University of Utah); Eric M. Reiman, MD (Banner Alzheimer’s Institute); Kewei Chen, PhD (Banner Alzheimer’s Institute); Chet Mathis, MD (University of Pittsburgh); Susan Landau, PhD (UC Berkeley); Nigel J. Cairns, PhD, MRCPath (Washington University St. Louis); Erin Householder (Washington University St. Louis); Lisa Taylor Reinwald, BA, HTL (Washington University St. Louis); Virginia Lee, PhD, MBA (UPenn School of Medicine); Magdalena Korecka, PhD (UPenn School of Medicine); Michal Figurski, PhD (UPenn School of Medicine); Karen Crawford (USC); Scott Neu, PhD (USC); Tatiana M. Foroud, PhD (Indiana University); Steven Potkin, MD UC (UC Irvine); Li Shen, PhD (Indiana University); Faber Kelley, MS, CCRC (Indiana University); Sungeun Kim, PhD (Indiana University); Kwangsik Nho, PhD (Indiana University); Zaven Kachaturian, PhD (Khachaturian, Radebaugh & Associates, Inc and Alzheimer’s Association’s Ronald and Nancy Reagan’s Research Institute); Richard Frank, MD, PhD (General Electric); Peter J. Snyder, PhD (Brown University); Susan Molchan, PhD (National Institute on Aging/ National Institutes of Health); Jeffrey Kaye, MD (Oregon Health and Science University); Joseph Quinn, MD (Oregon Health and Science University); Betty Lind, BS (Oregon Health and Science University); Raina Carter, BA (Oregon Health and Science University); Sara Dolen, BS (Oregon Health and Science University); Lon S. Schneider, MD (University of Southern California); Sonia Pawluczyk, MD (University of Southern California); Mauricio Beccera, BS (University of Southern California); Liberty Teodoro, RN (University of Southern California); Bryan M. Spann, DO, PhD (University of Southern California); James Brewer, MD, PhD (University of California San Diego); Helen Vanderswag, RN (University of California San Diego); Adam Fleisher, MD (University of California San Diego); Judith L. Heidebrink, MD, MS (University of Michigan); Joanne L. Lord, LPN, BA, CCRC (University of Michigan); Ronald Petersen, MD, PhD (Mayo Clinic, Rochester); Sara S. Mason, RN (Mayo Clinic, Rochester); Colleen S. Albers, RN (Mayo Clinic, Rochester); David Knopman, MD (Mayo Clinic, Rochester); Kris Johnson, RN (Mayo Clinic, Rochester); Rachelle S. Doody, MD, PhD (Baylor College of Medicine); Javier Villanueva Meyer, MD (Baylor College of Medicine); Munir Chowdhury, MBBS, MS (Baylor College of Medicine); Susan Rountree, MD (Baylor College of Medicine); Mimi Dang, MD (Baylor College of Medicine); Yaakov Stern, PhD (Columbia University Medical Center); Lawrence S. Honig, MD, PhD (Columbia University Medical Center); Karen L. Bell, MD (Columbia University Medical Center); Beau Ances, MD (Washington University, St. Louis); John C. Morris, MD (Washington University, St. Louis); Maria Carroll, RN, MSN (Washington University, St. Louis); Sue Leon, RN, MSN (Washington University, St. Louis); Erin Householder, MS, CCRP (Washington University, St. Louis); Mark A. Mintun, MD (Washington University, St. Louis); Stacy Schneider, APRN, BC, GNP (Washington University, St. Louis); Angela Oliver, RN, BSN, MSG; Daniel Marson, JD, PhD (University of Alabama Birmingham); Randall Griffith, PhD, ABPP (University of Alabama Birmingham); David Clark, MD (University of Alabama Birmingham); David Geldmacher, MD (University of Alabama Birmingham); John Brockington, MD (University of Alabama Birmingham); Erik Roberson, MD (University of Alabama Birmingham); Hillel Grossman, MD (Mount Sinai School of Medicine); Effie Mitsis, PhD (Mount Sinai School of Medicine); Leyla deToledo-Morrell, PhD (Rush University Medical Center); Raj C. Shah, MD (Rush University Medical Center); Ranjan Duara, MD (Wien Center); Daniel Varon, MD (Wien Center); Maria T. Greig, HP (Wien Center); Peggy Roberts, CNA (Wien Center); Marilyn Albert, PhD (Johns Hopkins University); Chiadi Onyike, MD (Johns Hopkins University); Daniel D’Agostino II, BS (Johns Hopkins University); Stephanie Kielb, BS (Johns Hopkins University); James E. Galvin, MD, MPH (New York University); Dana M. Pogorelec (New York University); Brittany Cerbone (New York University); Christina A. Michel (New York University); Henry Rusinek, PhD (New York University); Mony J de Leon, EdD (New York University); Lidia Glodzik, MD, PhD (New York University); Susan De Santi, PhD (New York University); P. Murali Doraiswamy, MD (Duke University Medical Center); Jeffrey R. Petrella, MD (Duke University Medical Center); Terence Z. Wong, MD (Duke University Medical Center); Steven E. Arnold, MD (University of Pennsylvania); Jason H. Karlawish, MD (University of Pennsylvania); David Wolk, MD (University of Pennsylvania); Charles D. Smith, MD (University of Kentucky); Greg Jicha, MD (University of Kentucky); Peter Hardy, PhD (University of Kentucky); Partha Sinha, PhD (University of Kentucky); Elizabeth Oates, MD (University of Kentucky); Gary Conrad, MD (University of Kentucky); Oscar L. Lopez, MD (University of Pittsburgh); MaryAnn Oakley, MA (University of Pittsburgh); Donna M. Simpson, CRNP, MPH (University of Pittsburgh); Anton P. Porsteinsson, MD (University of Rochester Medical Center); Bonnie S. Goldstein, MS, NP (University of Rochester Medical Center); Kim Martin, RN (University of Rochester Medical Center); Kelly M. Makino, BS (University of Rochester Medical Center); M. Saleem Ismail, MD (University of Rochester Medical Center); Connie Brand, RN (University of Rochester Medical Center); Ruth A. Mulnard, DNSc, RN, FAAN (University of California, Irvine); Gaby Thai, MD (University of California, Irvine); Catherine Mc Adams Ortiz, MSN, RN, A/GNP (University of California, Irvine); Kyle Womack, MD (University of Texas Southwestern Medical School); Dana Mathews, MD, PhD (University of Texas Southwestern Medical School); Mary Quiceno, MD (University of Texas Southwestern Medical School); Ramon Diaz Arrastia, MD, PhD (University of Texas Southwestern Medical School); Richard King, MD (University of Texas Southwestern Medical School); Myron Weiner, MD (University of Texas Southwestern Medical School); Kristen Martin Cook, MA (University of Texas Southwestern Medical School); Michael DeVous, PhD (University of Texas Southwestern Medical School); Allan I. Levey, MD, PhD (Emory University); James J. Lah, MD, PhD (Emory University); Janet S. Cellar, DNP, PMHCNS BC (Emory University); Jeffrey M. Burns, MD (University of Kansas, Medical Center); Heather S. Anderson, MD (University of Kansas, Medical Center); Russell H. Swerdlow, MD (University of Kansas, Medical Center); Liana Apostolova, MD (University of California, Los Angeles); Kathleen Tingus, PhD (University of California, Los Angeles); Ellen Woo, PhD (University of California, Los Angeles); Daniel H.S. Silverman, MD, PhD (University of California, Los Angeles); Po H. Lu, PsyD (University of California, Los Angeles); George Bartzokis, MD (University of California, Los Angeles); Neill R Graff Radford, MBBCH, FRCP (London) (Mayo Clinic, Jacksonville); Francine Parfitt, MSH, CCRC (Mayo Clinic, Jacksonville); Tracy Kendall, BA, CCRP (Mayo Clinic, Jacksonville); Heather Johnson, MLS, CCRP (Mayo Clinic, Jacksonville); Martin R. Farlow, MD (Indiana University); Ann Marie Hake, MD (Indiana University); Brandy R. Matthews, MD (Indiana University); Scott Herring, RN, CCRC (Indiana University); Cynthia Hunt, BS, CCRP (Indiana University); Christopher H. van Dyck, MD (Yale University School of Medicine); Richard E. Carson, PhD (Yale University School of Medicine); Martha G. MacAvoy, PhD (Yale University School of Medicine); Howard Chertkow, MD (McGill Univ., Montreal Jewish General Hospital); Howard Bergman, MD (McGill Univ., Montreal Jewish General Hospital); Chris Hosein, Med (McGill Univ., Montreal Jewish General Hospital); Sandra Black, MD, FRCPC (Sunnybrook Health Sciences, Ontario); Dr Bojana Stefanovic (Sunnybrook Health Sciences, Ontario); Curtis Caldwell, PhD (Sunnybrook Health Sciences, Ontario); Ging Yuek Robin Hsiung, MD, MHSc, FRCPC (U.B.C. Clinic for AD & Related Disorders); Howard Feldman, MD, FRCPC (U.B.C. Clinic for AD & Related Disorders); Benita Mudge, BS (U.B.C. Clinic for AD & Related Disorders); Michele Assaly, MA Past (U.B.C. Clinic for AD & Related Disorders); Andrew Kertesz, MD (Cognitive Neurology St. Joseph’s, Ontario); John Rogers, MD (Cognitive Neurology St. Joseph’s, Ontario); Dick Trost, PhD (Cognitive Neurology St. Joseph’s, Ontario); Charles Bernick, MD (Cleveland Clinic Lou Ruvo Center for Brain Health); Donna Munic, PhD (Cleveland Clinic Lou Ruvo Center for Brain Health); Diana Kerwin, MD (Northwestern University); Marek Marsel Mesulam, MD (Northwestern University); Kristine Lipowski, BA (Northwestern University); Chuang Kuo Wu, MD, PhD (Northwestern University); Nancy Johnson, PhD (Northwestern University); Carl Sadowsky, MD (Premiere Research Inst (Palm Beach Neurology)); Walter Martinez, MD (Premiere Research Inst (Palm Beach Neurology)); Teresa Villena, MD (Premiere Research Inst (Palm Beach Neurology)); Raymond Scott Turner, MD, PhD (Georgetown University Medical Center); Kathleen Johnson, NP (Georgetown University Medical Center); Brigid Reynolds, NP (Georgetown University Medical Center); Reisa A. Sperling, MD (Brigham and Women’s Hospital); Keith A. Johnson, MD (Brigham and Women’s Hospital); Gad Marshall, MD (Brigham and Women’s Hospital); Meghan Frey (Brigham and Women’s Hospital); Jerome Yesavage, MD (Stanford University); Joy L. Taylor, PhD (Stanford University); Barton Lane, MD (Stanford University); Allyson Rosen, PhD (Stanford University); Jared Tinklenberg, MD (Stanford University); Marwan N. Sabbagh, MD (Banner Sun Health Research Institute); Christine M. Belden, PsyD (Banner Sun Health Research Institute); Sandra A. Jacobson, MD (Banner Sun Health Research Institute); Sherye A. Sirrel, MS (Banner Sun Health Research Institute); Neil Kowall, MD (Boston University); Ronald Killiany, PhD (Boston University); Andrew E. Budson, MD (Boston University); Alexander Norbash, MD (Boston University); Patricia Lynn Johnson, BA (Boston University); Thomas O. Obisesan, MD, MPH (Howard University); Saba Wolday, MSc (Howard University); Joanne Allard, PhD (Howard University); Alan Lerner, MD (Case Western Reserve University); Paula Ogrocki, PhD (Case Western Reserve University); Leon Hudson, MPH (Case Western Reserve University); Evan Fletcher, PhD (University of California, Davis Sacramento); Owen Carmichael, PhD (University of California, Davis Sacramento); John Olichney, MD (University of California, Davis Sacramento); Charles DeCarli, MD (University of California, Davis Sacramento); Smita Kittur, MD (Neurological Care of CNY); Michael Borrie, MB ChB (Parkwood Hospital); T Y Lee, PhD (Parkwood Hospital); Dr Rob Bartha, PhD (Parkwood Hospital); Sterling Johnson, PhD (University of Wisconsin); Sanjay Asthana, MD (University of Wisconsin); Cynthia M. Carlsson, MD (University of Wisconsin); Steven G. Potkin, MD (University of California, Irvine BIC); Adrian Preda, MD (University of California, Irvine BIC); Dana Nguyen, PhD (University of California, Irvine BIC); Pierre Tariot, MD (Banner Alzheimer’s Institute); Adam Fleisher, MD (Banner Alzheimer’s Institute); Stephanie Reeder, BA (Banner Alzheimer’s Institute); Vernice Bates, MD (Dent Neurologic Institute); Horacio Capote, MD (Dent Neurologic Institute); Michelle Rainka, PharmD, CCRP (Dent Neurologic Institute); Douglas W. Scharre, MD (Ohio State University); Maria Kataki, MD, PhD (Ohio State University); Anahita Adeli, MD (Ohio State University); Earl A. Zimmerman, MD (Albany Medical College); Dzintra Celmins, MD (Albany Medical College); Alice D. Brown, FNP (Albany Medical College); Godfrey D. Pearlson, MD (Hartford Hosp, Olin Neuropsychiatry Research Center); Karen Blank, MD (Hartford Hosp, Olin Neuropsychiatry Research Center); Karen Anderson, RN (Hartford Hosp, Olin Neuropsychiatry Research Center); Robert B. Santulli, MD (Dartmouth Hitchcock Medical Center); Tamar J. Kitzmiller (Dartmouth Hitchcock Medical Center); Eben S. Schwartz, PhD (Dartmouth Hitchcock Medical Center); Kaycee M. Sink, MD, MAS (Wake Forest University Health Sciences); Jeff D. Williamson, MD, MHS (Wake Forest University Health Sciences); Pradeep Garg, PhD (Wake Forest University Health Sciences); Franklin Watkins, MD (Wake Forest University Health Sciences); Brian R. Ott, MD (Rhode Island Hospital); Henry Querfurth, MD (Rhode Island Hospital); Geoffrey Tremont, PhD (Rhode Island Hospital); Stephen Salloway, MD, MS (Butler Hospital); Paul Malloy, PhD (Butler Hospital); Stephen Correia, PhD (Butler Hospital); Howard J. Rosen, MD (UC San Francisco); Bruce L. Miller, MD (UC San Francisco); Jacobo Mintzer, MD, MBA (Medical University South Carolina); Kenneth Spicer, MD, PhD (Medical University South Carolina); David Bachman, MD (Medical University South Carolina); Elizabether Finger, MD (St. Joseph’s Health Care); Stephen Pasternak, MD (St. Joseph’s Health Care); Irina Rachinsky, MD (St. Joseph’s Health Care); John Rogers, MD (St. Joseph’s Health Care); Andrew Kertesz, MD (St. Joseph’s Health Care); Dick Drost, MD (St. Joseph’s Health Care); Nunzio Pomara, MD (Nathan Kline Institute); Raymundo Hernando, MD (Nathan Kline Institute); Antero Sarrael, MD (Nathan Kline Institute); Susan K. Schultz, MD (University of Iowa College of Medicine, Iowa City); Laura L. Boles Ponto, PhD (University of Iowa College of Medicine, Iowa City); Hyungsub Shim, MD (University of Iowa College of Medicine, Iowa City); Karen Elizabeth Smith, RN (University of Iowa College of Medicine, Iowa City); Norman Relkin, MD, PhD (Cornell University); Gloria Chaing, MD (Cornell University); Lisa Raudin, PhD (Cornell University); Amanda Smith, MD (University of South Floriday: USF Health Byrd Alzheimer’s Institute); Kristin Fargher, MD (University of South Floriday: USF Health Byrd Alzheimer’s Institute); Balebail Ashok Raj, MD (University of South Floriday: USF Health Byrd Alzheimer’s Institute)

